# Recombination-aware phylogenomics unravels the complex divergence of hybridizing species

**DOI:** 10.1101/485904

**Authors:** Gang Li, Henrique V. Figueiró, Eduardo Eizirik, William J. Murphy

## Abstract

Current phylogenomic approaches implicitly assume that the predominant phylogenetic signal within a genome reflects the true evolutionary history of organisms, without assessing the confounding effects of gene flow that result in a mosaic of phylogenetic signals that interact with recombinational variation. Here we tested the validity of this assumption with a recombination-aware analysis of whole genome sequences from 27 species of the cat family. We found that the prevailing phylogenetic signal within the autosomes is not always representative of speciation history, due to ancient hybridization throughout felid evolution. Instead, phylogenetic signal was concentrated within large, conserved X-chromosome recombination deserts that exhibited recurrent patterns of strong genetic differentiation and selective sweeps across mammalian orders. By contrast, regions of high recombination were enriched for signatures of ancient gene flow, and these sequences inflated crown-lineage divergence times by ~40%. We conclude that standard phylogenomic approaches to infer the Tree of Life may be highly misleading without considering the genomic partitioning of phylogenetic signal relative to recombination rate, and its interplay with historical hybridization.

## Introduction

Efforts to resolve the Tree of Life have been empowered by a major shift towards whole genome phylogenetic inference, or phylogenomics (Delsuc et al. 2005; McCormack et al. 2012; Jarvis et al. 2014; Lamichhaney et al. 2015; Prum et al. 2015; Figueiró et al. 2017; Irisarri et al. 2018). Phylogenomic studies typically analyze thousands of orthologous loci or whole genome sequence (wgs) data simultaneously, using concatenation or coalescent methods. This majority rule approach has long been regarded as the most successful way to mitigate the distorting influences of evolutionary noise at individual loci, by assuming that the correct genealogical signal (the ‘species tree’) will emerge with an ever-increasing amount of data, although this assumption is disputed (Romiguier et al. 2013). Further, interspecific gene flow is increasingly recognized as an important driver of genomic variation across the Tree of Life (Nadeau et al. 2013; Cahill et al. 2015; Good et al. 2015; Lamichhaney et al. 2015; Nater et al. 2015; Li, Davis, et al. 2016). As a result, signatures of ancient branching events can be overwritten and effectively erased from chromosomal segments, notably in regions with high rates of meiotic recombination where haplotype blocks are smaller, leading to more localized effect of positive or background selection. A phylogenomic study of *Anopheles* mosquitoes provided a vivid illustration of this scenario in which autosomal gene trees were homogenized by rampant gene flow, misleading standard phylogenomic inference (Fontaine et al. 2015). The correct species tree signal was instead enriched within the smaller X chromosome, consistent with the large X-effect, which states that the X chromosome is enriched for genetic elements with large effects on reducing hybrid reproductive fitness (Presgraves 2008; Larson et al. 2017; Presgraves 2018). In that case, the species tree was determined by demonstrating that nodes on the X chromosome phylogeny were significantly older than their alternative counterparts in the most frequent autosomal topology. Furthermore, the inferred species tree was enriched within a region of the X chromosome known to harbor reproductive isolation loci. Whether this surprising finding in mosquitos was exceptional for that group of insects, or is typical of other taxonomic lineages, is unknown.

Given the demonstrable impact that gene flow may have on genome-wide patterns of phylogenetic variation, and the contrasting patterns of genetic differentiation between autosomes (less) and sex chromosomes (greater) that are predicted by the large X-effect (Presgraves 2018), we hypothesized that applying standard majority rule phylogenomic approaches may be highly misleading when applied to taxa with complex speciation histories. Here we tested this hypothesis by analyzing the genomes of 27 species of the cat family (Felidae), a mammalian radiation characterized by notoriously difficult phylogenetic problems (Johnson et al. 2006; Li, Davis, et al. 2016). Previous studies have consistently partitioned the 41 currently recognized felid species into eight clades that radiated rapidly during the mid-late Miocene (Johnson et al. 2006) (Fig. 1). However, phylogenies constructed from separate mitogenome, autosomal, and sex chromosome partitions have displayed strongly supported differences within and between the eight clades, which may be driven by complex effects of sex-biased dispersal, gene flow, demography, and/or selection (Johnson et al. 2006; Luo et al. 2014; Li, Davis, et al. 2016). Indeed, studies utilizing nuclear SNP markers have demonstrated that many discordant phylogeographic patterns within felids inferred from mitochondrial DNA data were likely due to interspecific gene flow (Fig. 1) (Davis et al. 2010; Li, Davis, et al. 2016; Figueiró et al. 2017). Likewise, the phylogeny derived from the paternally inherited Y chromosome was similar in many respects to the autosomal tree, but was also shown to be influenced by sex-biased dispersal and ancient gene flow (Luo et al. 2014; Li, Davis, et al. 2016). Biogeographic reconstructions of the early felid radiation revealed that the ancestors of all eight extant clades evolved in Eurasia, most likely in sympatry or parapatry, before dispersing across the globe (Li, Davis, et al. 2016). Taken together, previous studies suggest a scenario for the evolution of extant felid clades that is consistent with a history of divergence with gene flow.

**Fig. 1.**
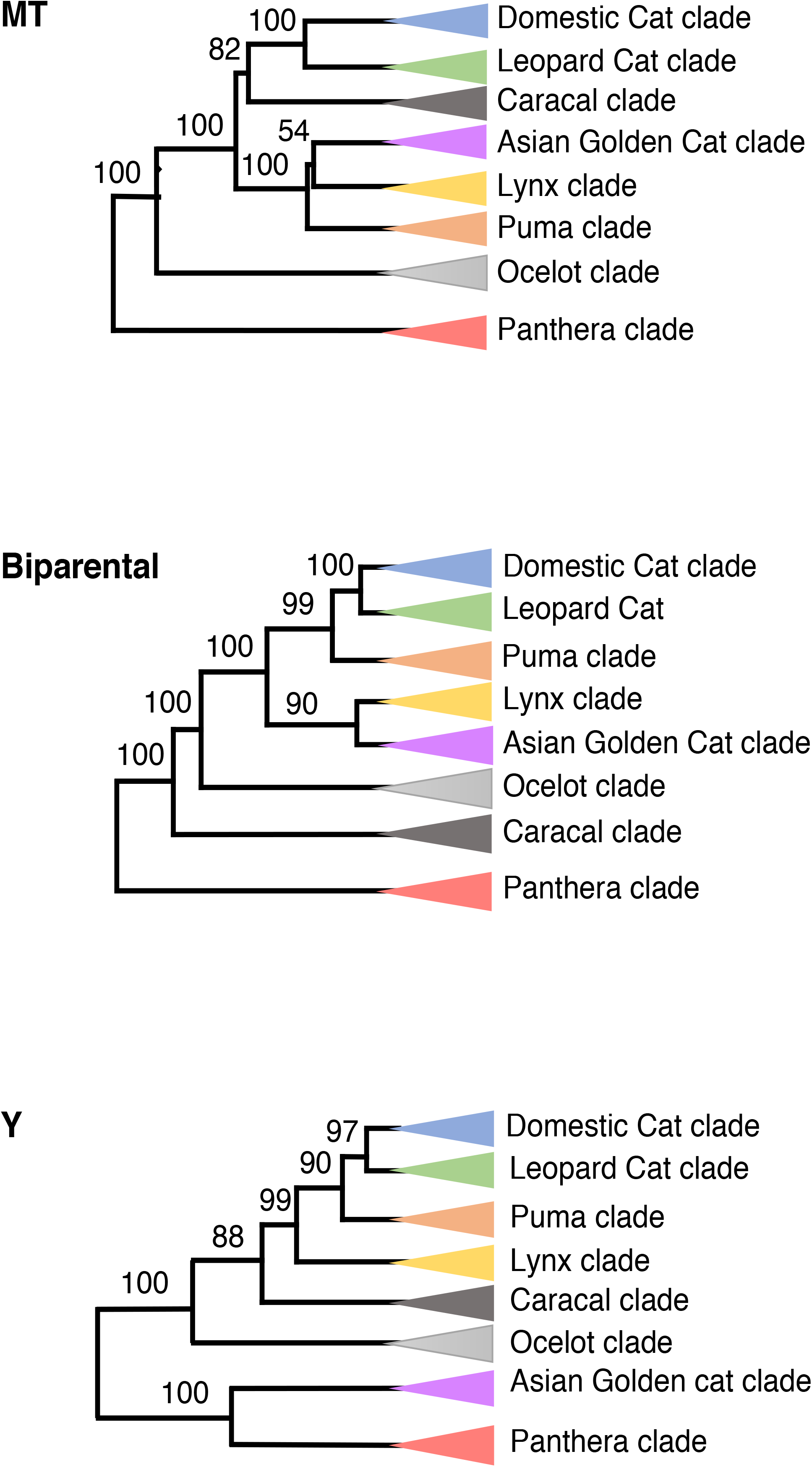
Previous hypotheses of felid phylogenetic relationships based on 1) maternally inherited mitogenome sequences (Li, Davis, et al. 2016), 2) biparentally inherited SNP markers (Li, Davis, et al. 2016), and 3) paternally inherited Y chromosome sequences (Johnson et al. 2006; Luo et al. 2014).

An important genetic parameter that influences assessments of gene flow, incomplete lineage sorting (ILS), and genetic diversity is the local rate of recombination (Ortíz-Barrientos et al. 2002; Hobolth et al. 2011; Nachman and Payseur 2012; Mailund et al. 2014; Burri et al. 2015; Payseur and Rieseberg 2016). This rate strongly influences how genealogical histories are distributed across chromosomes, as non-species trees are more likely to persist within regions with high recombination rates because deleterious alleles become effectively unlinked from neutral or positively selected variants (Brandvain et al. 2014; Schumer et al. 2018). However, recombination rate is generally not considered when selecting phylogenetic markers in spite of its predicted influence on phylogenetic discordance and estimates of divergence times across the genome (Schierup and Hein 2000; Ortíz-Barrientos et al. 2002; Posada and Crandall 2002; Lemey and Posada 2009). Specifically, few empirical studies have examined the direct impact of recombination rate on phylogenetic inference within rapidly diverging, speciose clades with histories of admixture, likely due to the rarity of recombination maps for the vast majority of organisms (but see (Pease and Hahn 2013)).

To more thoroughly understand the interplay between local recombination rates, inferred interspecies gene flow and phylogenomic resolution in the cat family, we analyzed the distribution of phylogenetic signal across the genomes of 27 felid species in the context of recombination rates inferred from high-resolution linkage maps for the domestic cat (Menotti-Raymond et al. 1999; Menotti-Raymond et al. 2003; Li, Hillier, et al. 2016). We found that phylogenomic signal is strongly influenced by ancient admixture, that it is partitioned within the nuclear genome by local recombination rates, and that the inferred species branching events are enriched on the X chromosome. Molecular dating results were also strongly influenced by ancient gene flow, leading to overestimation of divergence times from regions of high recombination that are enriched for multiple, discordant phylogenies. Our results offer whole-genome evidence that diversification in the presence of gene flow can distort phylogenetic signal throughout the majority of the genome, potentially leaving the most accurate depiction of ancient branching events within only a small fraction of the nuclear genome.

## Results

We generated new whole-genome sequences for 14 felid species, and combined them with published genomes for 13 other felids, producing an alignment for 27 felids that spanned 1.5-Gb, or 57% of the domestic cat euchromatic reference assembly (Table S1). To identify regional variation in phylogenetic signal, we partitioned the 18 autosomes and the X chromosome (chrX) into 23,707 non-overlapping windows of 100 kilobases (kb) and performed maximum likelihood (ML) phylogenetic inference on each window. We also evaluated smaller window sizes (50-kb), which produced similar results (Fig. S1). We also separately explored the phylogenetic signal in the autosomes and chrX, with each partition divided into regions of low versus high recombination rate inferred from high-resolution linkage maps (Li, Hillier, et al. 2016). Given the robust support for the monophyly of the eight main felid clades (Johnson et al. 2006; Walters-Conte et al. 2014; Li, Davis, et al. 2016), we present results from analyses performed in two stages. First, we present genome-wide analyses of topological frequency and divergence time variation to identify the most probable branching order within the six clades for which we had sequence data from at least three species. We reasoned that within-clade analyses (i.e. assessing recently diverged species) would minimize the impact of homoplasy and putative ancient (interclade) gene flow on observed patterns of variation and estimated divergence times (Posada 2000; Lemey and Posada 2009). We then followed up with an analysis of interclade relationships, selecting two species from each of the eight clades (with the exception of the Caracal clade, represented only by the serval), spanning the oldest crown node of each clade. For the latter analysis, we predicted that the impacts of a longer history of gene flow and recombination between lineages would produce higher levels of apparent homoplasy, leading to phylogenetic distortion and age inflation for each clade’s crown node.

### Intraclade phylogeny

Maximum likelihood estimation of phylogeny for autosomal windows confirmed a highly supported tree for each of the six felid clades for which there were at least three sampled species (Fig. 2A, Fig. S1). These results corroborated relationships observed in earlier phylogenetic studies that combined autosomal and sex chromosome sequences into a single supermatrix (Johnson et al. 2006). However, in half of the assessed felid clades, the frequency-based rank order of the top three topologies inferred from low-recombining regions of chrX (LRchrX) was strikingly different from the autosomal signal (Fig. 2A, Fig. S1). Furthermore, the distribution of the most frequently observed chrX trees was non-randomly distributed, with an alternative topology enriched within one or more multi-megabase spans of extremely reduced recombination (Fig. 2B). In such a case, which of the alternative topologies is most likely to be the species tree?

**Fig. 2.**
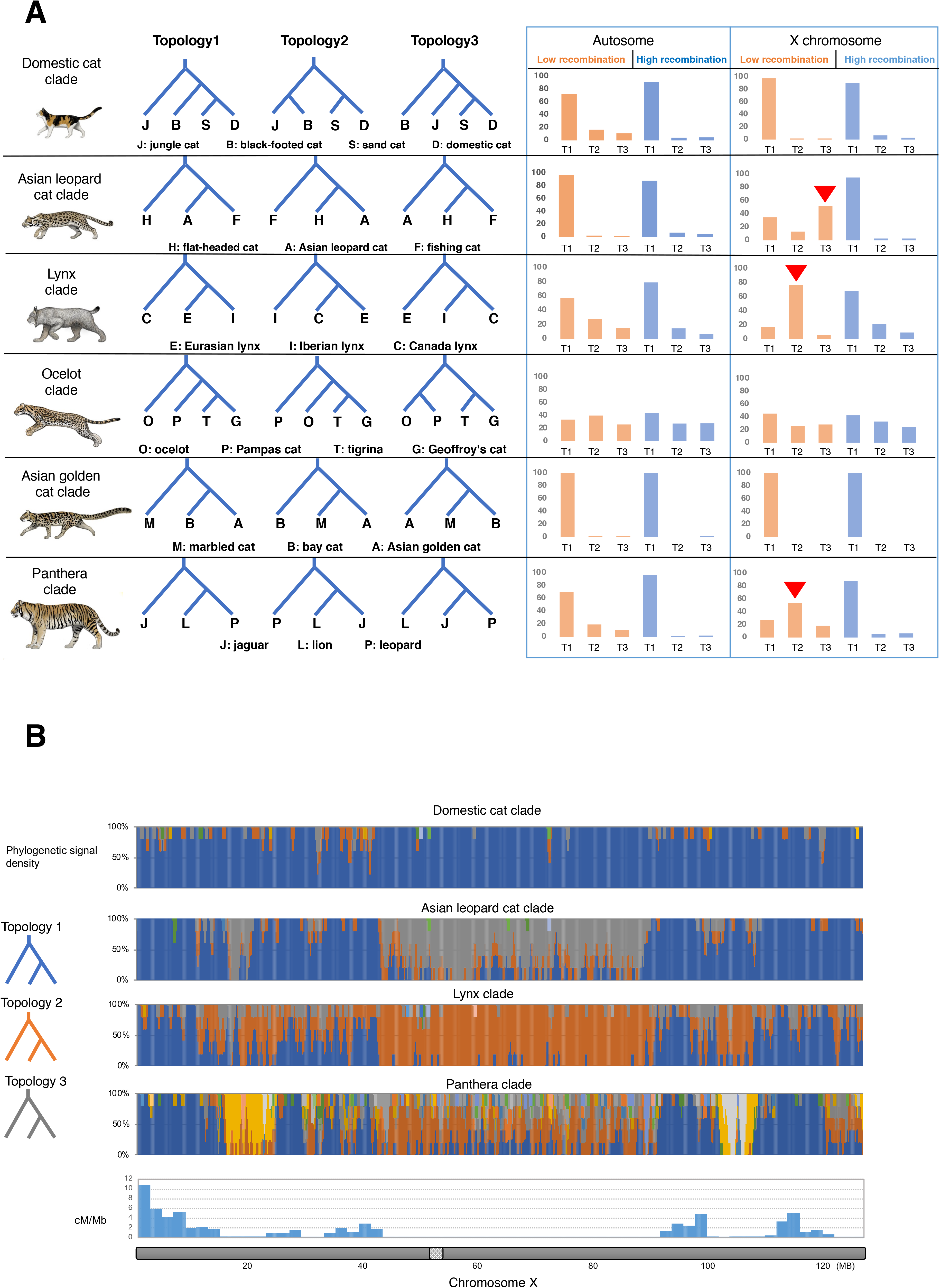
A. Left. The top three most frequently recovered phylogenies for six felid lineages, inferred from window-based maximum likelihood supermatrix analysis of the 1.5-Gb whole-genome alignment. Right. The frequency of each topology within autosomes and chrX, based on partitioning into low (<0.5 cM/Mb) and high (>2 cM/Mb) recombination rates (Li, Hillier, et al. 2016). Red arrows indicate three lineages where the most frequent topology in the LRchrX was not the same as the most frequent genome-wide topology. B. Distribution of different topologies on chrX, and their relationship to recombination rate (bottom).

The large X-effect predicts that the original branching history should be enriched on chrX, and previous studies indicate that low-recombining regions are more likely to retain the species tree because selection is more effective at removing weakly deleterious alleles (Nachman and Payseur 2012; Pease and Hahn 2013). This feature would enhance the probability of removing introgressed genomic blocks, since they are likely to contain alleles that are deleterious in their new genetic background. By contrast, non-species trees resulting from hybridization will tend to be enriched within regions displaying high recombination rates, because introgressed deleterious alleles are more efficiently unlinked from neutral or positively selected variants (Brandvain et al. 2014; Schumer et al. 2018). This would facilitate the incorporation of genomic blocks containing the latter two types of introgressed alleles in the descendant population. An additional criterion for determining the original branching events is the prediction that non-species trees with alternative relationships that are younger than their species-tree counterparts would likely be derived from post-speciation gene flow between non-sister taxa (Joly et al. 2009). In contrast, non-species trees that result from ILS should present divergence times at focal nodes (i.e. those defining competing relationships) that are older than those in the species tree. The third (and perhaps most commonly applied) criterion is the frequency of alternative trees relative to the inferred species tree, with ILS expected to generate non-species trees that are less prevalent than (or at most as prevalent as) the species tree. However, if post-speciation gene flow is the primary driver of genealogical discordance, the frequency of non-species trees could be considerably higher than the original branching signal (Fontaine et al. 2015).

Given these alternative predictions, and considering that age would be a key criterion to distinguish alternative scenarios (Joly et al. 2009; Fontaine et al. 2015), we estimated divergence times for the two competing hypotheses for each of the three felid lineages in Figure 2 which showed a marked genealogical disparity between the autosomes and chrX. We selected time estimated under a relaxed molecular clock (Drummond et al. 2006), using a standardized calibration, because raw sequence divergence levels may lead to overestimation of divergence time due to site-specific and/or lineage-specific rate heterogeneity. If the topology that predominated within LRchrX had a divergence time that was significantly younger than the most frequent tree in the remainder of the genome, this would be consistent with a scenario of post-speciation gene flow. In contrast, if the topology enriched within the LRchrX was older than the alternative topology, it would likely reflect a remnant of the original branching event.

In the Asian leopard cat and Lynx clades, the mean divergence time of the focal node for the most common topology in LRchrX (trees 3 and 2, respectively) was significantly older than that of the most common genome-wide topology (Figs. 3A & B), suggesting that the LRchrX tree reflected the ancient species branching relationships. Similarly, in other felid clades (except genus *Panthera*, Fig. 3C), we found that the inferred species tree was enriched within the LRchrX relative to high-recombination regions (Table S2). On the autosomes, trees supporting the inferred species branching history for the Asian leopard cat and Lynx clades were significantly less frequent (Figure 2), but still possessed the oldest divergence times (Figures 3A&B). Windows containing these trees also had significantly lower recombination rates than those supporting the most common topology (tree 1) (Table 1). These results support our inference that the tree enriched within LRchrX represents the correct species branching relationships for these two clades, and that species trees in general persist within regions of reduced recombination.

**Table 1.** Recombination rate (θ) differences between windows supporting the putative species tree^(ST)^ and the most common genome-wide tree in Fig. 3.

| | Mean $\theta$ | No. Windows | *P*-value<sup>1</sup> |
| --- | --- | --- | --- |
| Asian leopard cat clade |  |  |  |
| Topology 1-AUTO | 1.72 | 20,645 | < 0.001 |
| Topology 3<sup>ST</sup>-AUTO | 0.98 | 612 |  |
| Topology 1-chrX | 1.78 | 764 | < 0.001 |
| Topology 3<sup>ST</sup>-chrX | 0.12 | 361 |  |
| Lynx clade |  |  |  |
| Topology 1-AUTO | 1.85 | 14,449 | < 0.001 |
| Topology 2<sup>ST</sup>-AUTO | 1.39 | 5,242 |  |
| Topology 1-chrX | 2.28 | 455 | < 0.001 |
| Topology 2<sup>ST</sup>-chrX | 0.56 | 496 |  |
| Panthera clade |  |  |  |
| Topology 1-AUTO | 1.94 | 14,864 | < 0.001 |
| Topology 2<sup>ST</sup>-AUTO | 1.11 | 1,874 |  |
| Topology 1-chrX | 2.25 | 518 | < 0.001 |
| Topology 2<sup>ST</sup>-chrX | 0.23 | 293 |  |

<sup>1</sup>Mann-Whitney U test comparing the $\theta$ distribution of two topologies.

**Fig. 3.**
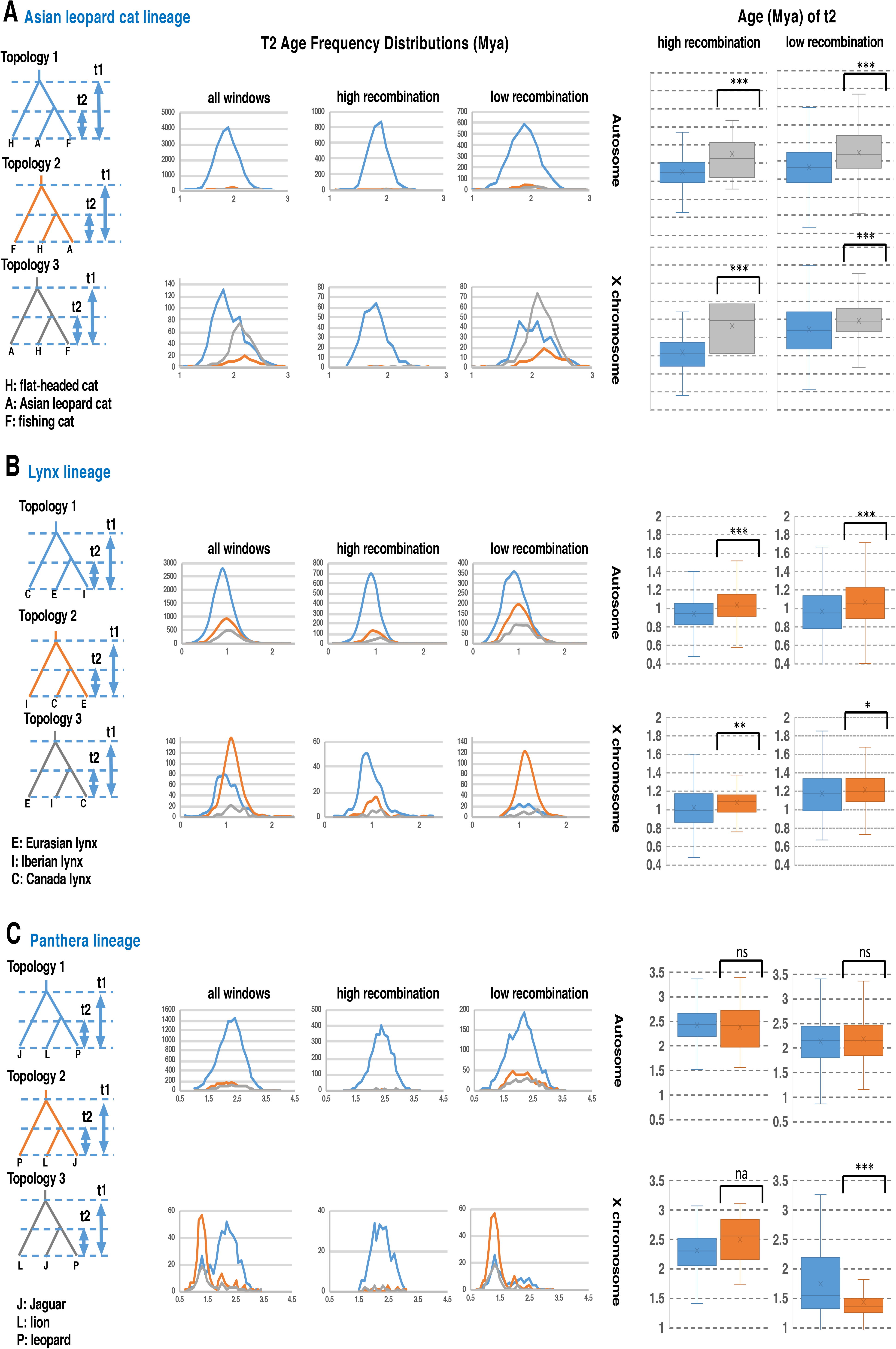
Left: Frequency distribution (*y*-axis) of window-based nodal estimates for divergence time 2 (t2) (*x*-axis) for the three lineages with discrepancies between autosomal and chrX divergence times in Fig. 1. Right: Divergence time estimates for the closest sister-species pair (time 2) found in the two most frequent topologies. In the Asian leopard cat (**A**) and lynx (**B**) lineages, the most frequent topology in LRchrX (gray or orange, respectively) has significantly older divergence times than those estimated for topology 1 (blue) (Mann-Whitney U test, ***P<0.001, **P<0.01, *P<0.05, ns=not significant, na=not analyzed due to too few windows). For the *Panthera* genus (**C**), divergence time estimates from LRchrX are consistent with recent hybridization between lion and jaguar, followed by selection-driven introgression (Figueiró et al. 2017).

The exceptional result for *Panthera* may have several explanations. In our previous studies of introgression within this genus (Li, Davis, et al. 2016; Figueiró et al. 2017) we demonstrated that divergence times for non-species trees were significantly reduced within the largest recombination cold spot on chrX due to selection-driven admixture, and we identified candidate loci that may have driven this sweep. Given this result, we inferred at the time that trees showing lion+jaguar as sister species were not the species tree, even though this particular topology was actually enriched within low-recombining regions throughout the genome (Figueiró et al. 2017), in contrast to theoretical expectations. However, it is possible that the lion+jaguar topology is the actual species tree, in spite of being less frequent in the genome. Indeed, windows supporting this topology have lower recombination rates on both the autosomes and chrX (Table 1). Such a result could be explained if there was post-speciation gene flow between sister species, resulting in much lower divergence times in low-recombining regions of chrX (Fig. 3C) in regions previously characterized by selective sweeps. In addition, it is noteworthy that *Panthera* tree topologies within the largest recombination cold spot on chrX are very compressed, raising the possibility that one or more selective sweeps occurred in the common ancestor of *Panthera* and/or throughout the divergence of the genus, leading to the consistently reduced divergence times we observe in this region (Figure 3C) (Nachman and Payseur 2012; O’Fallon 2013).

### Interclade phylogeny

Next, we explored the effect of recombination rate and mode of inheritance on the branching relationships of the eight main felid clades (Figure 1). When we divided chrX into low-versus high-recombination regions, ML trees from the high-recombination regions of chrX were identical to that of the autosomal partitions (separate or combined) and possessed very high bootstrap support (Fig. 4A). By contrast, the LRchrX tree relationships were also strongly supported, and differed from the chrX high-recombination region trees by both topology (i.e., the position of Asian golden cat and ocelot lineages) and most notably, branch lengths (Fig. 4A), indicating the signatures of two highly supported alternative phylogenetic histories for the cat family that were partitioned by recombination rates. The four most common topologies on chrX made up >95% of the chromosome (Fig. 4B). Interestingly, 99.7% of windows that supported Tree 1 were confined to the two largest recombination coldspots, whereas Trees 3 and 4 were largely restricted to regions characterized by high recombination rate, notably the pseudoautosomal region. Tree 2 was restricted to transitional regions between the lowest and highest recombination rates. Therefore, similar to our findings within the different felid clades, the LRchrX possessed a very distinct phylogenetic signal from high recombining regions on the same chromosome, as well as the autosomes.

**Fig. 4.**
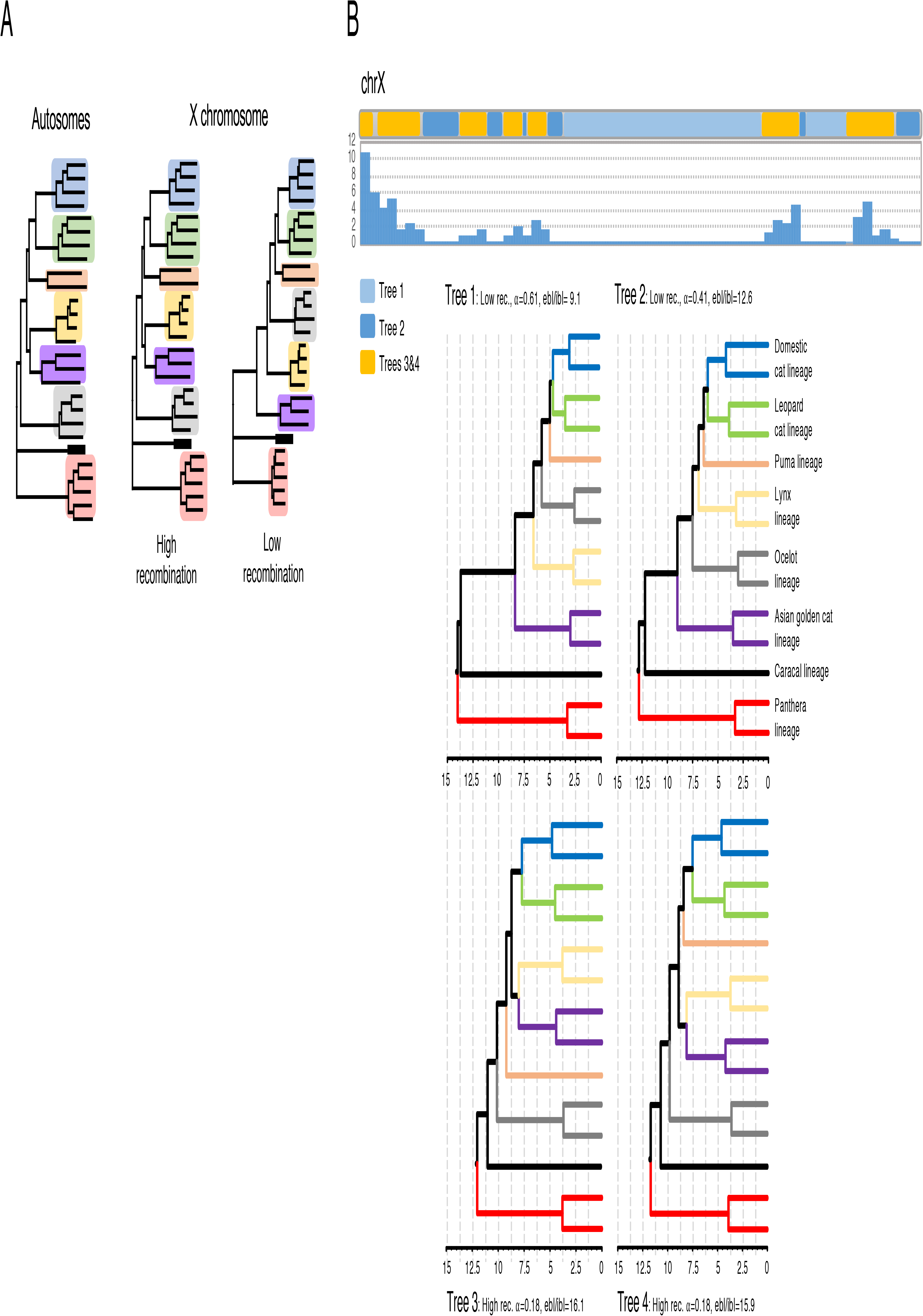
The influence of chrX recombination rate (*y*-axis, cM/Mb) on topology and divergence times, estimated in 100-kb windows along the chromosome. A. Trees are shown for autosomes (combined) and the chrX low and high recombination rates. Clade coloring follows Figure 1. B. Top: distribution of trees based on chrX recombination rate. Blocks were color-coded per topology, and smoothed in 5-Mb blocks. Bottom: Two measures that are distorted by hybridization and recombination are the ratio of external/internal branch length in a given tree (ebl/ibl) and α, the shape parameter of the gamma distribution. Timetrees show the impact of branch length distortion on divergence time estimation.

### Recombination influences phylogeny and divergence time estimation

Interspecific gene flow has predictable distorting effects on phylogeny (Posada and Crandall 2002; Lemey and Posada 2009), and this information can be used to assist in determining the most likely interclade branching events and divergence times. Frequent genetic exchange may produce star-like phylogenies when adjacent sites with different histories are combined. This has the specific effect of compressing internal branch lengths (ibl) relative to external branches (ebl), with this distortion being proportional to the topological distance between the hybridizing lineages (Schierup and Hein 2000; Lemey and Posada 2009). As a result, simulation studies have shown that phylogenies constructed from segments of the genome with larger numbers of sites reflecting discordant evolutionary histories are predicted to possess lower values of the alpha (shape) parameter of the gamma distribution of rate variation among sites, as multiple changes will be required at some sites to compensate for apparent homoplasy (Lemey and Posada 2009). Therefore, we calculated both alpha and the ebl/ibl ratio for the four most common topologies on chrX (Fig. 4B), and showed that both parameters were strongly influenced by the local recombination rate. ML analyses of high-recombination windows (Trees 3 and 4) produced more star-like trees, with 1.7-fold higher ratios of external to internal branch lengths, and threefold lower alpha values than trees produced from recombination cold spots (Fig. 4A&B), confirming predictions based on simulations (Schierup and Hein 2000; Lemey and Posada 2009). Our findings accord well with previous gene-centric phylogenomic studies which suggested that genes with high GC content (which has been reported to correlate with high rates of meiotic recombination) were associated with increased phylogenetic conflict (Romiguier et al. 2013; Shen et al. 2016).

We then assessed the effects of local recombination rates along chrX on the distribution of divergence time estimates for crown felid lineages (Fig. 4B, Table S3). The windows with the highest recombination rates on chrX contain a higher proportion of mixed phylogenetic signals from interspecific gene flow, and accordingly produced divergence times as much as 69% older than estimates derived from the LRchrX (tree 1). This fits the prediction that many apparent homoplasies are introduced on branches affected by gene flow (Lemey and Posada 2009). Previous phylogenetic studies comparing molecular divergence times for felid lineages with estimates from the fossil record noted that the latter underestimated, on average, the first occurrence of crown felid clades by 76% (Johnson et al. 2006; Li, Davis, et al. 2016). This assumes that these previous molecular dates represent unbiased estimates of divergence time. By comparison, our revised divergence times from the LRchrX mitigate the distorting effects of recombinant sequences and remove much of the previously implied ghost lineage history for different felid clades. Our revised molecular divergence time estimates from LRchrX are thus in much stronger agreement with known felid fossils from each clade (Table S3) (Johnson et al. 2006).

### Tests for gene flow and genomic distribution of admixture signatures

While incomplete lineage sorting certainly underpins some weakly-resolved aspects of discordant branching patterns within and between clades, it is unlikely to be responsible for the striking differences we observed between the autosomes and chrX. Previous analyses that applied moderate-density SNP variation to resolve felid phylogenetic relationships provided strong evidence for ancient, interspecies gene flow within several of the main felid clades (Li, Davis, et al. 2016). Here, we applied a combination of four- and five-taxon *D*-statistic tests to our whole-genome sequence data, assuming the LRchrX tree as the species phylogeny, to reexamine these results at much higher resolution, and to assess the possibility that gene flow between the ancestors of the eight modern felid clades was the proximate cause of previous difficulties resolving these early branching events. Our results provide significant evidence for ancient hybridization within and between nearly all clades of the Felidae (Fig. 5, Table S4). The observed patterns of ancient interspecific gene flow within each of the felid clades support and extend earlier findings from nuclear SNP datasets (Walters-Conte et al. 2014; Li, Davis, et al. 2016). For example, the intra-clade gene flow evident between the Asian leopard cat and both Fishing cat and Rusty spotted cat reconciles the discordant phylogeographic patterns of mitochondrial haplotypes observed in prior studies (Walters-Conte et al. 2014; Li, Davis, et al. 2016) that likely result from mitochondrial capture. Similarly, previous whole genome-based demographic evidence supported a scenario of historic gene flow between the Iberian lynx and Eurasian lynx (Abascal et al. 2016), consistent with our *D* statistic-based inferences of strong gene flow between the two species.

**Fig. 5.**
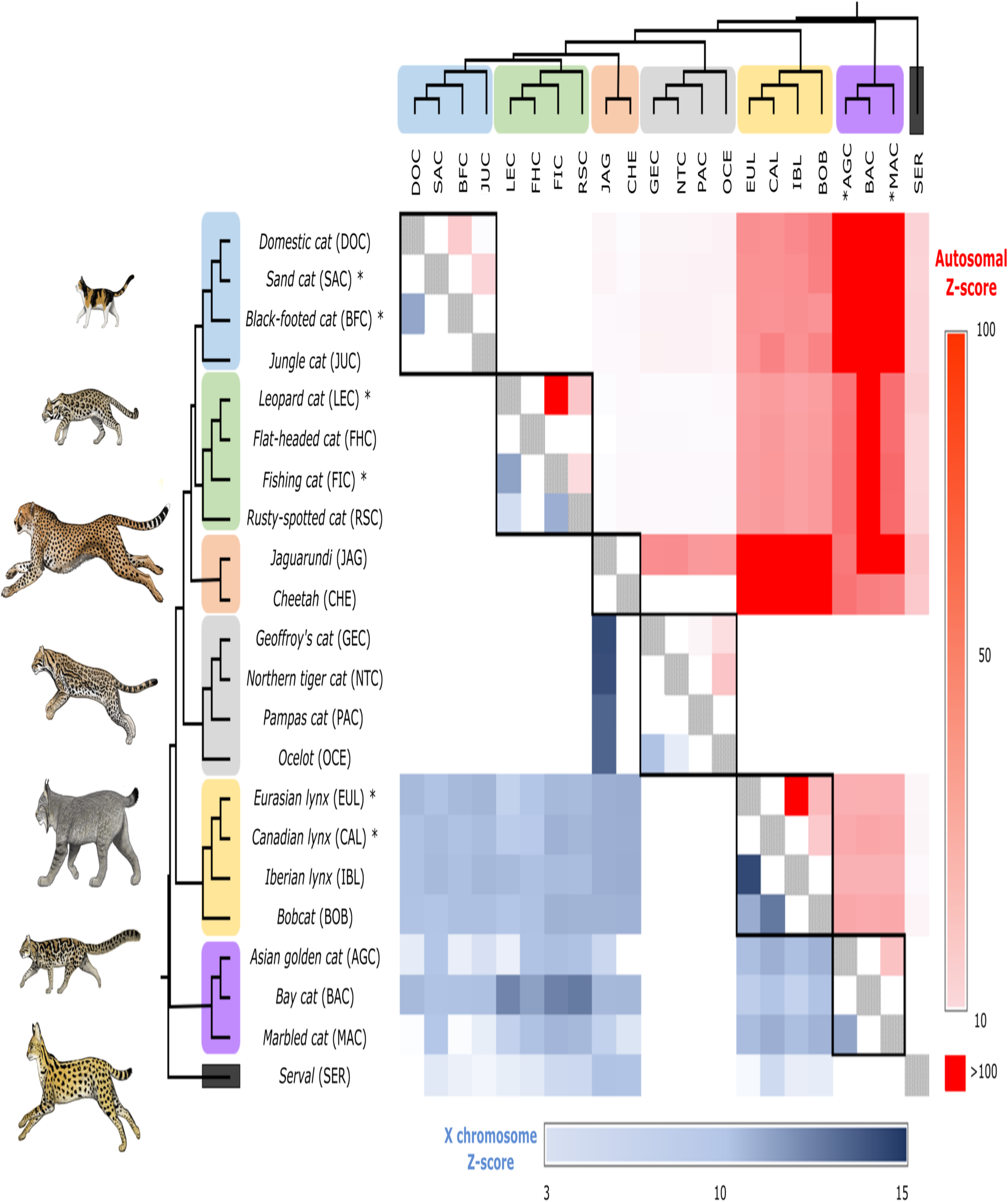
Matrix of pairwise *Z*-scores for the *D*-statistic estimated from the ABBA-BABA tests performed on all possible combinations of species trios within the non-pantherine felids, assuming the species tree inferred from chrX (Figure 4B) with *Panthera* as an outgroup. *D* statistics for *Panthera* have been previously reported (Figueiró et al. 2017) and are not shown. *Z* scores for the autosomes (upper, red shades) and the X chromosome (lower, blue shades) are shown separately. Intra-lineage scores are displayed inside black boxes along the diagonal. Asterisks indicate species that were included in the inter-lineage comparisons shown in Figure 6.

We also quantified and mapped inter-clade introgressed blocks using five-taxon comparisons performed with the software Dfoil, and observed striking patterns of ancient admixture, indicating that unique signatures of ancient introgression are consistently retained in each descendant species (Fig. 6, Figs. S2-S4). For example, we examined three different inferred episodes of ancient inter-clade admixture involving the Asian Golden Cat (AGC) clade, in one case connecting it to the Lynx clade, in another to the Domestic cat clade, and in the third to the Asian leopard cat clade (Fig. 6). In each case, we focused on the most widespread signatures of introgression, which connected the ancestor of the two extant species of the other clade and each of the sampled species of the AGC clade. Due to a limitation of the Dfoil method, we could not directly assess hybridization between the two ancestral branches (one from each clade), and thus indirectly investigated these episodes via the present-day signatures remaining in each of the two sampled species of the AGC clade. Remarkably, in all three cases, one of the species (marbled cat) consistently retains a larger number of introgressed segments, which are present across all chromosomes but tend to be concentrated in higher-recombination, telomeric regions. The other species (Asian golden cat) consistently retains fewer segments from each of the three ancient episodes, and these segments are more evenly-distributed across the genome (Fig. 6).

**Fig. 6.**
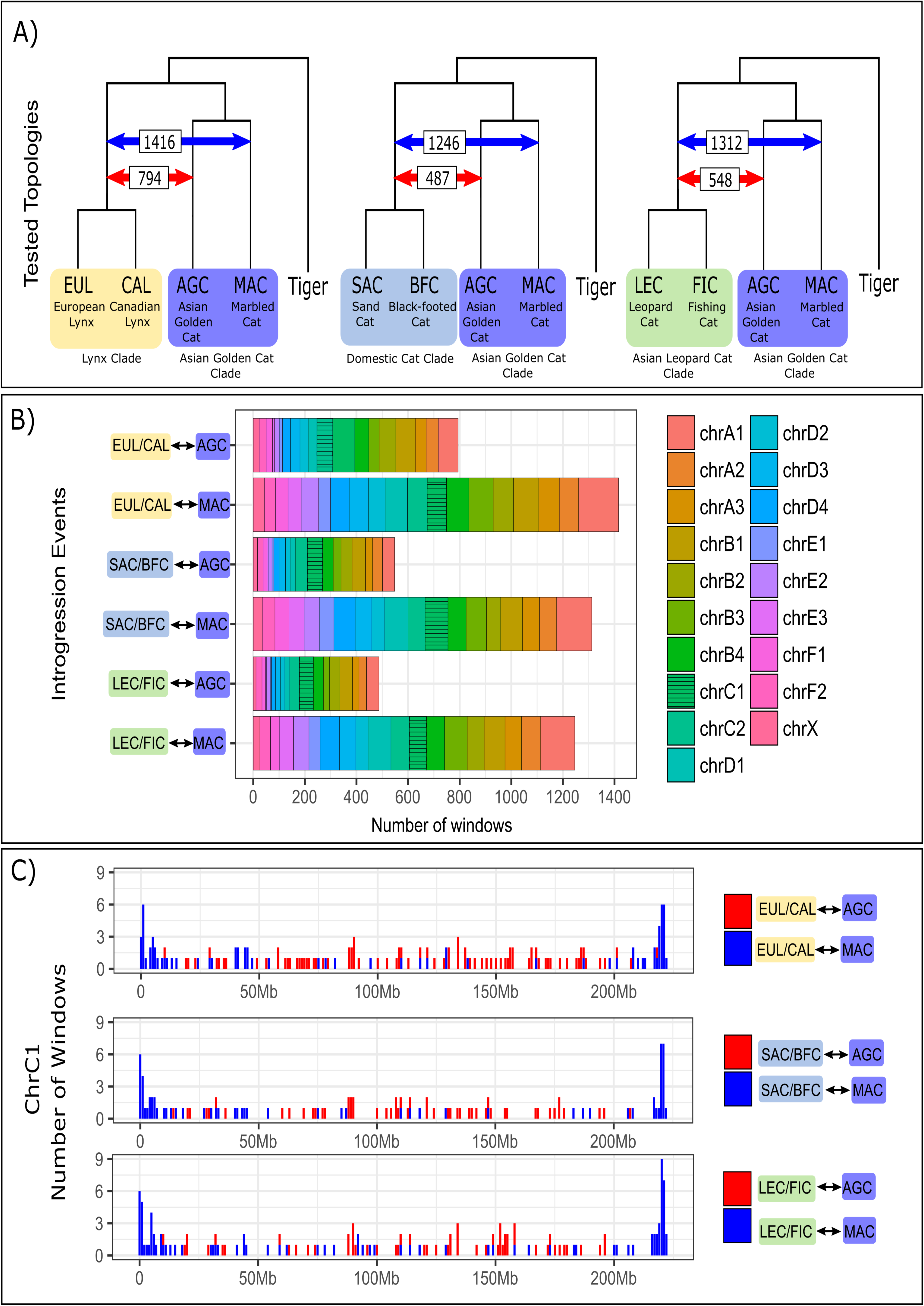
Quantification and chromosomal distribution of introgressed blocks in the genomes of two felid species (Asian golden cat and marbled cat), resulting from hybridization of their common ancestor with the progenitors of three different lineages of the Felidae. A) Red and Blue arrows show the estimated number of windows which support gene flow between two lineages. B) Chromosome distribution of windows with signatures of gene flow. C) Spatial distribution of introgressed blocks derived from the hybridization events shown are shown for chromosome C1 (see Figure S2-S4 for full results).

Such a consistent difference suggests that these two species have distinct genetic/evolutionary properties (e.g., demographic history and effective population size (*N*_e_)) that lead them to retain different signatures of ancient introgression (putatively from the same episodes, preceding their divergence from each other). To investigate this hypothesis, we performed PSMC analyses for these two species (Fig. S5), and observed distinct trajectories, with the Asian golden cat tending to have a larger historical and recent *N*_e_, relative to the marbled cat. Although this past demographic trend may not have had a direct influence on the observed difference in introgression signatures, it raises the hypothesis that *N*_e_ differences not only affect the immediate fitness consequences of hybridization (Schumer et al. 2018), but may also shape present-day signatures of ancient hybridization events through their effect on genetic drift (and thus retention of alternative haplotypes present in the population). If this hypothesis is affirmed by future assessments, it would imply an additional layer of complexity for phylogenomic inference, since the demographic history of present-day species (and its interplay with selection and recombination) would influence the magnitude and pattern of retention of alternative genealogies, potentially leading to considerably different results depending on the species that is sampled to represent a given clade.

## Discussion

Our results provide empirical support for the large X-effect within members of the cat family, where regions of suppressed recombination on chrX are enriched for the original species branching relationships and display greater differentiation between closely related species than high-recombination regions on the same chromosome. Previous studies have demonstrated that low-recombining regions should retain ancient branching events in the presence of gene flow, as linkage disequilibrium with genes contributing to reproductive isolation that accumulate within such regions would lead to removal of deleterious alleles introduced through hybridization (Nachman and Payseur 2012; Pease and Hahn 2013). Within the domestic cat genome, the multimegabase chrX recombination cold spots are by far the largest, and possess much lower levels of recombination than are observed on any autosomes (Li, Hillier, et al. 2016). While numerous empirical studies have associated genes contributing to reproductive isolation with chromosomal inversions that suppress recombination, within felids there are no such inversions that we could attribute to the large chrX recombination cold spots. So what other evolutionary mechanisms might be responsible for such a large region of significantly reduced recombination that retains species branching histories?

X chromosome collinearity is unique among eutherian mammal chromosomes in its exceptional conservation, and the gene order we observe across the massive recombination cold spots in cats is conserved between numerous eutherian mammals (Murphy et al. 1999; Raudsepp et al. 2004; O’Brien et al. 2006; Delgado et al. 2009; Ma et al. 2010). We compared the high-resolution recombination and physical maps available for human (Nagaraja et al. 1997), cat (Li, Hillier, et al. 2016) and pig (Ma et al. 2010) and observed a remarkable conservation of recombination rate profiles across chrX (Fig. 7) that span more than 180 million years of divergence (Meredith et al. 2011). While the distribution and pattern of recombination rates have been shown to persist at megabase levels between human and chimpanzees (Paigen and Petkov 2010), there is no precedent for such extreme conservation across deep timescales. Intriguingly, the largest 40 Mb recombination coldspot in the center of chrX preserves independent signals of putative selective sweeps, with nearly identical, orthologous breakpoints in *Panthera* and in pigs (Ai et al. 2015; Li, Davis, et al. 2016; Figueiró et al. 2017). This large region also largely overlaps syntenic segments in primates marked by selective sweeps, and extended regions of strong genetic differentiation between subspecies and species of rabbits and sheep with similar boundaries (Carneiro et al. 2010; Carneiro et al. 2014; Chen et al. 2018)(Dutheil et al. 2015; Nam et al. 2015). A previous study also found that the same recombination coldspot was among the most strongly differentiated genomic regions distinguishing domestic cats from their wild progenitors (Montague et al. 2014).

**Fig. 7.**
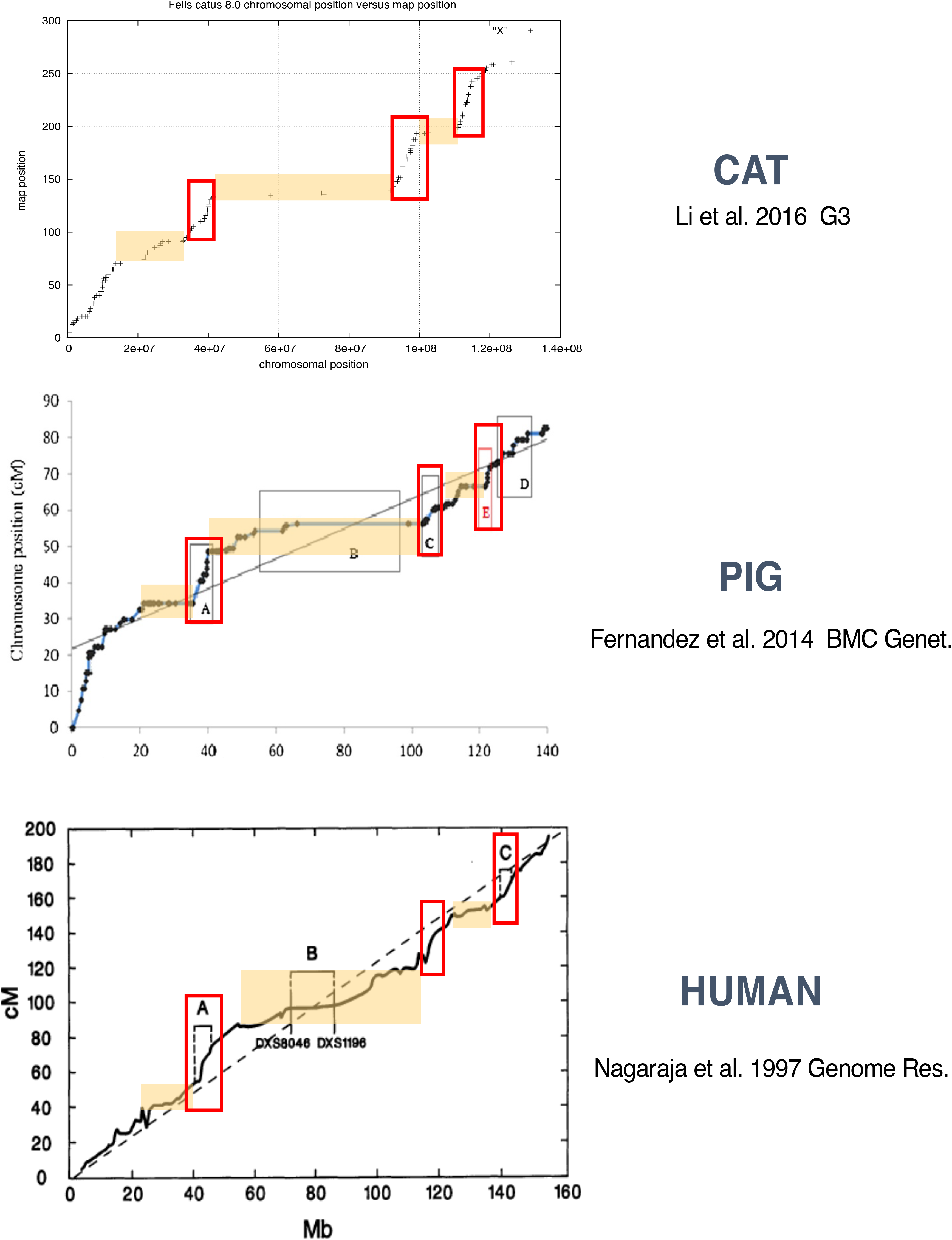
Remarkable conservation of gene order and recombination rate across eutherian mammal orders. Comparisons of recombination maps of chrX in cat (Li, Hillier, et al. 2016), pig (Fernández et al. 2014) and human (Nagaraja et al. 1997). Images are reproduced from the original publications (including lettered notations that refer to shifts in recombination rate). Red boxes highlight conserved, syntenic regions of elevated recombination rate, while yellow boxes highlight conserved syntenic regions of reduced recombination rate.

It would appear unlikely that the loci driving these signatures of genetic differentiation are identical, considering that more than 500 genes reside within the interval. However, there are notable functional features that are dispersed centrally within and adjacent to the conserved recombination coldspot that merit consideration as drivers of the recurrent patterns of strong genetic differentiation we observe across divergent mammalian orders. The X chromosome inactivation center (XIC), which encodes non-coding RNA loci that initiate and maintain female X chromosome silencing, lies within the center of the largest recombination coldspot. Extending outward from the XIC are multiple, dispersed loci that are known to physically interact to aid in the formation of the 3-dimensional structure of the inactive X chromosome (i.e., the Barr body) in female therian mammals (Deng et al. 2015; Darrow et al. 2016; Giorgetti et al. 2016; Jégu et al. 2017; Wang et al. 2018). It is noteworthy that these recombination coldspots are also flanked by conserved regions of extremely elevated recombination rate (Fig. 7). Collectively, these observations suggest that exceptional structural and functional constraints, such as those imparted by X chromosome inactivation, have preserved long-range gene order and recombination patterns over such large evolutionary distances (Pál and Hurst 2003), and may provide a genomic environment that is favorable for the accumulation of reproductive isolation loci. Future studies will be necessary to better understand the mechanistic and evolutionary basis for the conservation of chrX recombination patterns, and whether similar patterns of species tree retention within LRchrX are more taxonomically widespread.

## Conclusions

In summary, our findings show striking parallels to the phylogenomic study of mosquitoes that demonstrated that the majority of the species-tree signal was retained within a minority of the genome, the X chromosome (Fontaine et al. 2015). This phylogenomic pattern, likely a consequence of the large role of the X chromosome in reproductive isolation (Presgraves 2008; Presgraves 2018), calls into question the validity of majority rule approaches for phylogenetic inference in taxonomic lineages with sex chromosomes that exhibit interspecific gene flow. It is clear that X-linkage is alone not predictive of the species tree, as species-tree and non-species-tree topologies are non-randomly distributed on chrX, with the inferred species tree enriched only within regions with extremely reduced recombination. We have also demonstrated a strong topological bias that results from using phylogenies derived from high-recombination regions to estimate divergence time within clades with reticulate evolutionary histories. Phylogenies inferred from these regions are generally compressed, which artificially leads to a reduction of divergence times at the root and an inflation of divergence time for crown clades near the tips (Posada 2000; Schierup and Hein 2000). This raises the prospect that some described ‘star-like’ phylogenies might be artifacts resulting from combined effects of selecting phylogenetic markers from regions of elevated recombination rate that contained mixed histories derived from historical/contemporary gene flow. Accounting for this bias may play a central role in resolving conflicts between molecular divergence times and paleontological estimates for taxon originations (Meredith et al. 2011; dos Reis et al. 2012; Liu et al. 2017; Springer et al. 2017), thus contributing to a substantially more accurate reconstruction of the history of life.

## Materials and methods

### Sampling and sequencing

We isolated genomic DNA from cultured fibroblast cells or frozen tissue from 14 wild felid species: Fishing cat (*Prionailurus viverrinus*), Flat-headed cat (*Prionailurus planiceps*), Rusty-spotted cat (*Prionailurus rubiginosus*), Asian golden cat (*Catopuma temminckii*), Bay cat (*Catopuma badia*), Marbled cat (*Pardofelis marmorata*), Canada lynx (*Lynx canadensis*), Bobcat (*Lynx rufus*), Jaguarundi (*Herpailurus yagouaroundi*), Serval (*Leptailurus serval*), Pampas cat (*Leopardus colocola*), Ocelot (*Leopardus pardalis*), Northern Tigrina (*Leopardus tigrinus*), and Geoffroy’s cat (*Leopardus geoffroyi*). DNA was extracted with the Qiagen DNeasy kit. Standard Illumina fragment libraries (~300-bp average insert size) were prepared for each DNA sample and sequenced to ~30× genome-wide depth of coverage with 2×125-bp reads on the Illumina HiSeq4000 platform. We combined these new data with published genomes from 13 other felids: domestic cat, *Felis catus* (Montague et al. 2014); Sand cat, *Felis margarita* (Li, Davis, et al. 2016); Black-footed cat, *Felis nigripes* (Li, Davis, et al. 2016); Jungle cat, *Felis chaus* (Li, Davis, et al. 2016); Leopard cat, *Prionailurus bengalensis* (Li, Davis, et al. 2016); Eurasian lynx, *Lynx lynx* (Abascal et al. 2016); Iberian lynx, *Lynx pardinus* (Abascal et al. 2016); Cheetah, *Acinonyx jubatus* (Dobrynin et al. 2015); tiger, *Panthera tigris* (Cho et al. 2013); snow leopard, *Panthera uncia* (Cho et al. 2013), lion, *Panthera leo* (Cho et al. 2013); jaguar, *Panthera onca* (Figueiró et al. 2017); and leopard, *Panthera pardus* (Figueiró et al. 2017). These 27 species sample the main eight felid clades (Johnson et al. 2006), and comprise 77% of all currently recognized living cat species.

### Sequence QC and read mapping

All raw reads were trimmed using Trim Galore (http://www.bioinformatics.babraham.ac.uk/ projects/trim_galore). Trimmed and filtered reads were mapped to a repeatmasked version of the domestic cat genome (v8.0), which was assembled under the guidance of a high-resolution SNP array-based linkage map (Li, Hillier, et al. 2016). Sequence alignment was carried out using BWA with previously described mapping parameters (Li, Davis, et al. 2016). We applied the standard GATK pipeline (https://software.broadinstitute.org/gatk/) to remove PCR duplicates, perform local realignment, and calculate base quality score recalibration (BQSR).

### Genome Alignment and Phylogenomic Analysis

We used ANGSD (Korneliussen et al. 2014) to generate consensus pseudohaploid variant calls for each species, with minimum mapping quality set to 30, and minimum base quality set to 20. Segmental duplications and other highly repetitive regions evolve rapidly, make up as much as 5% of a mammalian genome, and collapse into single locations in short-read Illumina assemblies. Because these collapsed duplications can introduce errors into variant calling and grossly bias branch length estimation, we applied CNVnator (Abyzov et al. 2011) to identify and exclude all alignment regions with read depth greater than 150% or less than 50% of the genome-wide average in one or more species.

We evaluated the mapping quality of non-domestic felid sequence reads against the *Felis catus* v8 reference (Li, Hillier, et al. 2016), by comparison to variants called from alignment of the *de novo* assembly scaffolds/contigs. We used SOAPdenovo2 (Luo et al. 2012) to generate a primary *de novo* assembly (k-mer size of 31), then aligned all filtered contigs to the domestic cat repeatmasked reference genome. In the filtration step, all contigs <300-bp were removed. We used the whole genome aligner LAST (Kiełbasa et al. 2011) to perform pairwise alignments of the filtered assembly contigs.

We used RAxML (Stamatakis 2014) to perform maximum likelihood (ML) tree searches (GTR+GAMMA model of sequence evolution) and estimate the alpha parameter of the gamma distribution. Nodal support was estimated with 200 bootstrap replicates. We performed sliding window-based estimation of phylogeny and divergence times across the 27-species whole genome alignment. Alignment blocks were created and analyzed in non-overlapping, 100-kb contiguous windows across the whole genome alignment. We evaluated smaller (Fig. S1) and larger block sizes, and this had little effect on topology frequencies, therefore the 100-kb block size was used for all subsequent analyses. MCMCTree was used to calculate the relative divergence time for each node with known fossil calibrations (Johnson et al. 2006). Analyses were run for 100,000 generations, with burn-in set to 10,000 generations, and autocorrelated rates.

### *D* statistics

We applied methods that utilize patterns in DNA sequence variation to detect hybridization over evolutionary timescales, as implemented in ANGSD (Korneliussen et al. 2014). Specifically, we calculated *D* statistics and *Z*-scores with the ABBA/BABA method (Green et al. 2010), which identifies imbalances in alternative topology frequencies under a four-taxon scenario (three ingroup taxa and one outgroup) (Table S4). Pseudohaploid sequences were generated from each genome assembly. The statistical significance of the *Z*-score was assessed with a block jackknife test (5-Mb block size for autosomes and 10-Mb block size for chromosome X). In addition, we used D_FOIL_ (Pease and Hahn 2015) to estimate *D*-statistics with 5-taxon data sets (four ingroup taxa with symmetrical relationships and one outgroup) and 100-kb non-overlapping windows. We focused on taxon combinations that could test inter-lineage introgression episodes (hypothesized based on the results from the standard 4-taxon ABBA-BABA tests). We used the D_FOIL_ approach to quantify the introgressed blocks (number of 100-kb windows per 10-Mb segment) and to map their chromosomal locations (using the domestic cat assembly coordinates) (Figs S2-S4).

### Demographic History

We applied the pairwise sequentially Markovian coalescent (PSMC) (Li and Durbin 2011) method to infer the historical demography of two cat species (Asian golden cat and marbled cat) that exhibited distinct results in the D_FOIL_ analysis. To call diploid sequences, we generated *de novo* assemblies using SOAPdenovo2 (Luo et al. 2012) with kmer=31. Illumina sequences were mapped to their own *de novo* genome assembly using BWA with default parameter settings. SAMTOOLS was used to estimate average mapping coverage, and to call and filter nucleotide variants. Genomic regions with less than half or more than twice the average whole genome mapping depth were excluded from the final analysis. We applied a mutation rate of 1×10^−8^ and published generation times for both species (Sunquist and Sunquist 2017). The consistency of the PSMC results was assessed by performing 100 bootstrap replicates.

### Genome recombination rate

We used a high-density domestic cat linkage map (Li, Hillier, et al. 2016) to plot recombination rate variation (cM/Mb) across the genome in non-overlapping 500-Kb windows. We calculated the regional average recombination rate within sliding 2-Mb blocks.

## Acknowledgements

We would like to thank Al Roca, Nicole Foley, and Molly Schumer for helpful comments and discussion, and Stephen O’Brien for samples. This work was supported by the US National Science Foundation (DEB-1753760 to WJM), CAPES/Brazil (EE and HVF) and CNPq/Brazil (EE and HVF). The authors declare they have no competing interests. Whole genome sequence reads have been deposited under PRJNA407940 in the NCBI short read archive.

## References

Abascal F, Corvelo A, Cruz F, Villanueva-Cañas JL, Vlasova A, Marcet-Houben M, Martínez-Cruz B, Cheng JY, Prieto P, Quesada V, et al. 2016. Extreme genomic erosion after recurrent demographic bottlenecks in the highly endangered Iberian lynx. Genome Biol. 17:251.

Abyzov A, Urban AE, Snyder M, Gerstein M. 2011. CNVnator: an approach to discover, genotype, and characterize typical and atypical CNVs from family and population genome sequencing. Genome Res. 21:974–984.

Ai H, Fang X, Yang B, Huang Z, Chen H, Mao L, Zhang F, Zhang L, Cui L, He W, et al. 2015. Adaptation and possible ancient interspecies introgression in pigs identified by whole-genome sequencing. Nat. Genet. 47:217–225.

Brandvain Y, Kenney AM, Flagel L, Coop G, Sweigart AL. 2014. Speciation and introgression between Mimulus nasutus and Mimulus guttatus. PLoS Genet. 10:e1004410.

Burri R, Nater A, Kawakami T, Mugal CF, Olason PI, Smeds L, Suh A, Dutoit L, Bureš S, Garamszegi LZ, et al. 2015. Linked selection and recombination rate variation drive the evolution of the genomic landscape of differentiation across the speciation continuum of Ficedula flycatchers. Genome Res. 25:1656–1665.

Cahill JA, Stirling I, Kistler L, Salamzade R, Ersmark E, Fulton TL, Stiller M, Green RE, Shapiro B. 2015. Genomic evidence of geographically widespread effect of gene flow from polar bears into brown bears. Mol. Ecol. 24:1205–1217.

Carneiro M, Albert FW, Afonso S, Pereira RJ, Burbano H, Campos R, Melo-Ferreira J, Blanco-Aguiar JA, Villafuerte R, Nachman MW, et al. 2014. The genomic architecture of population divergence between subspecies of the European rabbit. PLoS Genet. 10:e1003519.

Carneiro M, Blanco-Aguiar JA, Villafuerte R, Ferrand N, Nachman MW. 2010. Speciation in the European rabbit (Oryctolagus cuniculus): islands of differentiation on the X chromosome and autosomes. Evolution 64:3443–3460.

Chen Z-H, Zhang M, Lv F-H, Ren X, Li W-R, Liu M-J, Nam K, Bruford MW, Li M-H. 2018. Contrasting Patterns of Genomic Diversity Reveal Accelerated Genetic Drift but Reduced Directional Selection on X-Chromosome in Wild and Domestic Sheep Species. Genome Biol. Evol. 10:1282–1297.

Cho YS, Hu L, Hou H, Lee H, Xu J, Kwon S, Oh S, Kim H-M, Jho S, Kim S, et al. 2013. The tiger genome and comparative analysis with lion and snow leopard genomes. Nat. Commun. 4:2433.

Darrow EM, Huntley MH, Dudchenko O, Stamenova EK, Durand NC, Sun Z, Huang S-C, Sanborn AL, Machol I, Shamim M, et al. 2016. Deletion of DXZ4 on the human inactive X chromosome alters higher-order genome architecture. Proc. Natl. Acad. Sci. U. S. A. 113:E4504–E4512.

Davis BW, Li G, Murphy WJ. 2010. Supermatrix and species tree methods resolve phylogenetic relationships within the big cats, Panthera (Carnivora: Felidae). Mol. Phylogenet. Evol. 56:64–76.

Delgado CLR, Waters PD, Gilbert C, Robinson TJ, Graves JAM. 2009. Physical mapping of the elephant X chromosome: conservation of gene order over 105 million years. Chromosome Res. 17:917–926.

Delsuc F, Brinkmann H, Philippe H. 2005. Phylogenomics and the reconstruction of the tree of life. Nat. Rev. Genet. 6:361.

Deng X, Ma W, Ramani V, Hill A, Yang F, Ay F, Berletch JB, Blau CA, Shendure J, Duan Z, et al. 2015. Bipartite structure of the inactive mouse X chromosome. Genome Biol. 16:152.

Dobrynin P, Liu S, Tamazian G, Xiong Z, Yurchenko AA, Krasheninnikova K, Kliver S, Schmidt-Küntzel A, Koepfli K-P, Johnson W, et al. 2015. Genomic legacy of the African cheetah, Acinonyx jubatus. Genome Biol. 16:277.

Drummond AJ, Ho SYW, Phillips MJ, Rambaut A. 2006. Relaxed phylogenetics and dating with confidence. PLoS Biol. 4:e88.

Dutheil JY, Munch K, Nam K, Mailund T, Schierup MH. 2015. Strong Selective Sweeps on the X Chromosome in the Human-Chimpanzee Ancestor Explain Its Low Divergence. PLoS Genet. 11:e1005451.

Fernández AI, Muñoz M, Alves E, Folch JM, Noguera JL, Enciso MP, Rodríguez MDC, Silió L. 2014. Recombination of the porcine X chromosome: a high density linkage map. BMC Genet. 15:148.

Figueiró HV, Li G, Trindade FJ, Assis J, Pais F, Fernandes G, Santos SHD, Hughes GM, Komissarov A, Antunes A, et al. 2017. Genome-wide signatures of complex introgression and adaptive evolution in the big cats. Sci Adv 3:e1700299.

Fontaine MC, Pease JB, Steele A, Waterhouse RM, Neafsey DE, Sharakhov IV, Jiang X, Hall AB, Catteruccia F, Kakani E, et al. 2015. Mosquito genomics. Extensive introgression in a malaria vector species complex revealed by phylogenomics. Science 347:1258524.

Giorgetti L, Lajoie BR, Carter AC, Attia M, Zhan Y, Xu J, Chen CJ, Kaplan N, Chang HY, Heard E, et al. 2016. Structural organization of the inactive X chromosome in the mouse. Nature 535:575–579.

Good JM, Vanderpool D, Keeble S, Bi K. 2015. Negligible nuclear introgression despite complete mitochondrial capture between two species of chipmunks. Evolution 69:1961–1972.

Green RE, Krause J, Briggs AW, Maricic T, Stenzel U, Kircher M, Patterson N, Li H, Zhai W, Fritz MH-Y, et al. 2010. A draft sequence of the Neandertal genome. Science 328:710–722.

Hobolth A, Dutheil JY, Hawks J, Schierup MH, Mailund T. 2011. Incomplete lineage sorting patterns among human, chimpanzee, and orangutan suggest recent orangutan speciation and widespread selection. Genome Res. 21:349–356.

Irisarri I, Singh P, Koblmüller S, Torres-Dowdall J, Henning F, Franchini P, Fischer C, Lemmon AR, Lemmon EM, Thallinger GG, et al. 2018. Phylogenomics uncovers early hybridization and adaptive loci shaping the radiation of Lake Tanganyika cichlid fishes. Nat. Commun. 9:3159.

Jarvis ED, Mirarab S, Aberer AJ, Li B, Houde P, Li C, Ho SYW, Faircloth BC, Nabholz B, Howard JT, et al. 2014. Whole-genome analyses resolve early branches in the tree of life of modern birds. Science 346:1320–1331.

Jégu T, Aeby E, Lee JT. 2017. The X chromosome in space. Nat. Rev. Genet. 18:377.

Johnson WE, Eizirik E, Pecon-Slattery J, Murphy WJ, Antunes A, Teeling E, O’Brien SJ. 2006. The late Miocene radiation of modern Felidae: a genetic assessment. Science 311:73–77.

Joly S, McLenachan PA, Lockhart PJ. 2009. A statistical approach for distinguishing hybridization and incomplete lineage sorting. Am. Nat. 174:E54–E70.

Kiełbasa SM, Wan R, Sato K, Horton P, Frith MC. 2011. Adaptive seeds tame genomic sequence comparison. Genome Res. 21:487–493.

Korneliussen TS, Albrechtsen A, Nielsen R. 2014. ANGSD: Analysis of Next Generation Sequencing Data. BMC Bioinformatics 15:356.

Lamichhaney S, Berglund J, Almén MS, Maqbool K, Grabherr M, Martinez-Barrio A, Promerová M, Rubin C-J, Wang C, Zamani N, et al. 2015. Evolution of Darwin’s finches and their beaks revealed by genome sequencing. Nature 518:371–375.

Larson EL, Keeble S, Vanderpool D, Dean MD, Good JM. 2017. The Composite Regulatory Basis of the Large X-Effect in Mouse Speciation. Mol. Biol. Evol. 34:282–295.

Lemey P, Posada D. 2009. Introduction to recombination detection. In: Lemey P, Salemi M, Vandamme A-M, Lemey P, Salemi M, Vandamme A-M, editors. The Phylogenetic Handbook. 2nd ed. Cambridge: Cambridge University Press. p. 493–518.

Li G, Davis BW, Eizirik E, Murphy WJ. 2016. Phylogenomic evidence for ancient hybridization in the genomes of living cats (Felidae). Genome Res. 26:1–11.

Li G, Hillier LW, Grahn RA, Zimin AV, David VA, Menotti-Raymond M, Middleton R, Hannah S, Hendrickson S, Makunin A, et al. 2016. A high-resolution SNP array-based linkage map anchors a new domestic cat draft genome assembly and provides detailed patterns of recombination. G3 6:1607–1616.

Li H, Durbin R. 2011. Inference of human population history from individual whole-genome sequences. Nature 475:493–496.

Liu L, Zhang J, Rheindt FE, Lei F, Qu Y, Wang Y, Zhang Y, Sullivan C, Nie W, Wang J, et al. 2017. Genomic evidence reveals a radiation of placental mammals uninterrupted by the KPg boundary. Proc. Natl. Acad. Sci. U. S. A. 114:E7282–E7290.

Luo R, Liu B, Xie Y, Li Z, Huang W, Yuan J, He G, Chen Y, Pan Q, Liu Y, et al. 2012. SOAPdenovo2: an empirically improved memory-efficient short-read de novo assembler. Gigascience 1:18.

Luo S-J, Zhang Y, Johnson WE, Miao L, Martelli P, Antunes A, Smith JLD, O’Brien SJ. 2014. Sympatric Asian felid phylogeography reveals a major Indochinese-Sundaic divergence. Mol. Ecol. 23:2072–2092.

Mailund T, Munch K, Schierup MH. 2014. Lineage sorting in apes. Annu. Rev. Genet. 48:519–535.

Ma J, Iannuccelli N, Duan Y, Huang W, Guo B, Riquet J, Huang L, Milan D. 2010. Recombinational landscape of porcine X chromosome and individual variation in female meiotic recombination associated with haplotypes of Chinese pigs. BMC Genomics 11:159.

McCormack JE, Faircloth BC, Crawford NG, Gowaty PA, Brumfield RT, Glenn TC. 2012. Ultraconserved elements are novel phylogenomic markers that resolve placental mammal phylogeny when combined with species-tree analysis. Genome Res. 22:746–754.

Menotti-Raymond M, David VA, Lyons LA, Schäffer AA, Tomlin JF, Hutton MK, O’Brien SJ. 1999. A genetic linkage map of microsatellites in the domestic cat (Felis catus). Genomics 57:9–23.

Menotti-Raymond M, David VA, Roelke ME, Chen ZQ, Menotti KA, Sun S, Schäffer AA, Tomlin JF, Agarwala R, O’Brien SJ, et al. 2003. Second-generation integrated genetic linkage/radiation hybrid maps of the domestic cat (Felis catus). J. Hered. 94:95–106.

Meredith RW, Janečka JE, Gatesy J, Ryder OA, Fisher CA, Teeling EC, Goodbla A, Eizirik E, Simão TLL, Stadler T, et al. 2011. Impacts of the Cretaceous Terrestrial Revolution and KPg extinction on mammal diversification. Science 334:521–524.

Montague MJ, Li G, Gandolfi B, Khan R, Aken BL, Searle SMJ, Minx P, Hillier LW, Koboldt DC, Davis BW, et al. 2014. Comparative analysis of the domestic cat genome reveals genetic signatures underlying feline biology and domestication. Proc. Natl. Acad. Sci. U. S. A. 111:17230–17235.

Murphy WJ, Sun S, Chen ZQ, Pecon-Slattery J, O’Brien SJ. 1999. Extensive conservation of sex chromosome organization between cat and human revealed by parallel radiation hybrid mapping. Genome Res. 9:1223–1230.

Nachman MW, Payseur BA. 2012. Recombination rate variation and speciation: theoretical predictions and empirical results from rabbits and mice. Philos. Trans. R. Soc. Lond. B Biol. Sci. 367:409–421.

Nadeau NJ, Martin SH, Kozak KM, Salazar C, Dasmahapatra KK, Davey JW, Baxter SW, Blaxter ML, Mallet J, Jiggins CD. 2013. Genome-wide patterns of divergence and gene flow across a butterfly radiation. Mol. Ecol. 22:814–826.

Nagaraja R, MacMillan S, Kere J, Jones C, Griffin S, Schmatz M, Terrell J, Shomaker M, Jermak C, Hott C, et al. 1997. X chromosome map at 75-kb STS resolution, revealing extremes of recombination and GC content. Genome Res. 7:210–222.

Nam K, Munch K, Hobolth A, Dutheil JY, Veeramah KR, Woerner AE, Hammer MF, Great Ape Genome Diversity Project, Mailund T, Schierup MH. 2015. Extreme selective sweeps independently targeted the X chromosomes of the great apes. Proc. Natl. Acad. Sci. U. S. A. 112:6413–6418.

Nater A, Burri R, Kawakami T, Smeds L, Ellegren H. 2015. Resolving Evolutionary Relationships in Closely Related Species with Whole-Genome Sequencing Data. Syst. Biol. 64:1000–1017.

O’Brien SJ, Menninger JC, Nash WG. 2006. Atlas of Mammalian Chromosomes. John Wiley & Sons

O’Fallon B. 2013. Purifying selection causes widespread distortions of genealogical structure on the human X chromosome. Genetics 194:485–492.

Ortíz-Barrientos D, Reiland J, Hey J, Noor MAF. 2002. Recombination and the divergence of hybridizing species. Genetica 116:167–178.

Paigen K, Petkov P. 2010. Mammalian recombination hot spots: properties, control and evolution. Nat. Rev. Genet. 11:221–233.

Pál C, Hurst LD. 2003. Evidence for co-evolution of gene order and recombination rate. Nat. Genet. 33:392–395.

Payseur BA, Rieseberg LH. 2016. A genomic perspective on hybridization and speciation. Mol. Ecol. 25:2337–2360.

Pease JB, Hahn MW. 2013. More accurate phylogenies inferred from low-recombination regions in the presence of incomplete lineage sorting. Evolution 67:2376–2384.

Pease JB, Hahn MW. 2015. Detection and Polarization of Introgression in a Five-Taxon Phylogeny. Syst. Biol. 64:651–662.

Posada D. 2000. How does recombination affect phylogeny estimation? Trends Ecol. Evol. 15:489–490.

Posada D, Crandall KA. 2002. The effect of recombination on the accuracy of phylogeny estimation. J. Mol. Evol. 54:396–402.

Presgraves DC. 2008. Sex chromosomes and speciation in Drosophila. Trends Genet. 24:336–343.

Presgraves DC. 2018. Evaluating genomic signatures of “the large X-effect” during complex speciation. Mol. Ecol. [Internet]. Available from: http://dx.doi.org/10.1111/mec.14777

Prum RO, Berv JS, Dornburg A, Field DJ, Townsend JP, Lemmon EM, Lemmon AR. 2015. A comprehensive phylogeny of birds (Aves) using targeted next-generation DNA sequencing. Nature 526:569–573.

Raudsepp T, Lee E-J, Kata SR, Brinkmeyer C, Mickelson JR, Skow LC, Womack JE, Chowdhary BP. 2004. Exceptional conservation of horse-human gene order on X chromosome revealed by high-resolution radiation hybrid mapping. Proc. Natl. Acad. Sci. U. S. A. 101:2386–2391.

dos Reis M, Inoue J, Hasegawa M, Asher RJ, Donoghue PCJ, Yang Z. 2012. Phylogenomic datasets provide both precision and accuracy in estimating the timescale of placental mammal phylogeny. Proc. Biol. Sci. 279:3491–3500.

Romiguier J, Ranwez V, Delsuc F, Galtier N, Douzery EJP. 2013. Less is more in mammalian phylogenomics: AT-rich genes minimize tree conflicts and unravel the root of placental mammals. Mol. Biol. Evol. 30:2134–2144.

Schierup MH, Hein J. 2000. Consequences of recombination on traditional phylogenetic analysis. Genetics 156:879–891.

Schumer M, Xu C, Powell D, Durvasula A, Skov L, Holland C, Sankararaman S, Andolfatto P, Rosenthal G, Przeworski M. 2018. Natural selection interacts with the local recombination rate to shape the evolution of hybrid genomes. Science [Internet]. Available from: http://dx.doi.org/10.1126/science.aar36842407

Shen X-X, Salichos L, Rokas A. 2016. A Genome-Scale Investigation of How Sequence, Function, and Tree-Based Gene Properties Influence Phylogenetic Inference. Genome Biol. Evol. 8:2565–2580.

Springer MS, Emerling CA, Meredith RW, Janečka JE, Eizirik E, Murphy WJ. 2017. Waking the undead: Implications of a soft explosive model for the timing of placental mammal diversification. Mol. Phylogenet. Evol. 106:86–102.

Stamatakis A. 2014. RAxML version 8: a tool for phylogenetic analysis and post-analysis of large phylogenies. Bioinformatics 30:1312–1313.

Sunquist M, Sunquist F. 2017. Wild Cats of the World. University of Chicago Press

Walters-Conte KB, Johnson DLE, Johnson WE, O’Brien SJ, Pecon-Slattery J. 2014. The dynamic proliferation of CanSINEs mirrors the complex evolution of Feliforms. BMC Evol. Biol. 14:137.

Wang C-Y, Jégu T, Chu H-P, Oh HJ, Lee JT. 2018. SMCHD1 Merges Chromosome Compartments and Assists Formation of Super-Structures on the Inactive X. Cell 174:406–421.e25.

